# Cryo-EM structure of plant ABCC1 reveals mode of arsenic transport

**DOI:** 10.64898/2026.11.16.767027

**Authors:** Madushani R.A. Wathukoralage, Alan T. Culbertson

## Abstract

Plant ATP-binding cassette transporter C1 (ABCC1) mediates the vacuolar sequestration of heavy metal-phytochelatin (PC) complexes for cytosolic detoxification, yet the structural mechanisms of substrate recognition and metal coordination remain uncharacterized. Here, we report functional assays and single-particle cryo-EM structures of maize ABCC1 (ZmABCC1) in its apo state and bound to an PC_2_-As(III) complex. Functional assays demonstrate that ZmABCC1 ATPase activity is sequentially increased by glutathione, PC_2_, and PC_3_ in the presence of As(III), supporting that longer PC chain lengths increase transport efficiency. The substrate-bound structure reveals the mode of substrate binding of the heavy metal complex, with the As(III) atom coordinated in a trigonal pyramidal geometry by three sulfur atoms from two PC_2_ ligands. Arg1200 plays a key role through nonpolar interaction with As(III) through its aliphatic side chain and an electrostatic interaction with the carboxyl group of PC_2_. These results define the molecular basis of PC-mediated heavy metal transport by plant ABCC1 which could be utilized to inform efforts to engineer crop stress resilience.

## Introduction

Heavy metals are naturally occurring elements, some of which are essential for biological processes, including iron (Fe), manganese (Mn), and zinc (Zn). However, several heavy metals such as lead (Pb), cadmium (Cd), mercury (Hg), and arsenic (As) lack biological roles and are toxic even at low concentrations (Ghori et al., 2019). Human activities in agriculture and mining have accelerated the environmental release of these toxic elements (Tchounwou et al., 2012). This environmental heavy metal contamination is a major problem for both human diseases and agricultural practices (Alengebawy et al., 2021; Witkowska et al., 2021). Further, unlike organic toxins, heavy metals cannot be degraded into non-toxic metabolites. Consequently, living organisms must rely on alternative detoxification mechanisms to limit toxicity, such as cellular export or intracellular sequestration (Ejaz et al., 2023).

Upon entering the eukaryotic cytosol, toxic metals are chelated by pathways involving thiol coordination by glutathione (GSH) or organic acid coordination by citrate and malate (Pearson and Cowan, 2021). The coordination mechanism depends on the preferred geometry and electronic properties of the metal ion. Arsenic, mercury, and cadmium are coordinated by thiol groups, whereas aluminum, iron, and calcium are coordinated by organic acids. Chelation itself provides initial cellular protection, but these complexes can be further detoxified by transport across the plasma membrane or sequestration into the vacuole. This transport is often mediated by ATP-binding cassette (ABC) transporters (Pearson and Cowan, 2021).

ABC transporters are membrane proteins that transport substrates across cellular membranes using energy derived from ATP binding and hydrolysis (Thomas and Tampe, 2020). Eukaryotic ABC proteins are divided into nine families (A through I), with families E and F performing non-transport functions, and families H and I occurring only in insects and plants, respectively (Huang et al., 2010; Chen et al., 2025). ABC transporters contain a minimum functional core structure consisting of two transmembrane domains (TMDs) and two nucleotide-binding domains (NBDs). Members of the ABCC subfamily often possess an additional N-terminal transmembrane domain, designated TMD0, which functions in protein localization or protein-protein interactions (Johnson and Chen, 2017; Shinde et al., 2025).

ABCC1, also known as Multidrug Resistance Protein 1 (MRP1), is a C-family ABC transporter involved in cytosolic detoxification (Cole and Deeley, 2006). Mammalian ABCC1 has been studied extensively due to its role in multidrug resistance during cancer chemotherapy (Cole, 2014). Functional homologs exist in plants (AtABCC1 and AtABCC2) and yeast (yeast cadmium factor 1, Ycf1), which display broad, overlapping substrate specificities compared with mammalian ABCC1 (Lu et al., 1997; Lu et al., 1998; Soong et al., 2025; Sun et al., 2025). These transporters mediate heavy metal detoxification and utilize amphiphilic anions, including glutathione (GS) or glucuronic acid conjugates (Leslie et al., 2004; Song et al., 2010; Park et al., 2012; Shukalek et al., 2016; Soong et al., 2025). The substrate binding mechanism is established, particularly for mammalian ABCC1, with several structures determined in complex with GS-conjugates or GSH co-bound with secondary molecules revealing key cationic residues that coordinate the anionic acid groups of GSH (Johnson and Chen, 2017; Sun et al., 2025). The structure of Arabidopsis ABCC2, a functional homolog of ABCC1, was recently determined bound to the GS-conjugated herbicide 2,4-dinitrophenyl-glutathione (Qiu et al., 2025). Structural studies of Ycf1 have demonstrated that linker phosphorylation regulates transporter activity (Bickers et al., 2021; Khandelwal et al., 2022) and has recently been determined bound to oxidized glutathione (GSSG) (Soong et al., 2025).

In addition to transporting small molecules either co-transported with GSH or as GS-conjugates, ABCC1 homologs from plants, mammals, and yeast mediate heavy metal detoxification (Li et al., 1997; Leslie et al., 2004; Song et al., 2010; Park et al., 2012; Shukalek et al., 2016). All three forms transport arsenic, cadmium, and mercury complexes, though individual homologs display distinct substrate preferences. For example, plant ABCC1 exhibits the highest transport rate for arsenic, whereas the yeast homolog exhibits highest transport activity for cadmium. Yeast and mammalian ABCC1 transport GS-complexed heavy metals in the form of As(GSH)_3_, Cd(GSH)_2_, Hg(GSH)_2_ (Shukalek et al., 2016). In contrast, plant ABCC1 utilizes phytochelatins (PCs), a polymerized form of GSH (Seregin and Kozhevnikova, 2023), and transports PC_2_ (Figure 1A) complexes with As(III) or Cd(II) at 2:1 and 1:1 stoichiometry, respectively (Song et al., 2010; Song et al., 2014). Despite multiple structures available for plant, mammalian, and yeast ABCC1 homologs, structural information for any of these transporters bound to a heavy metal complex remains absent.

**Figure 1.**
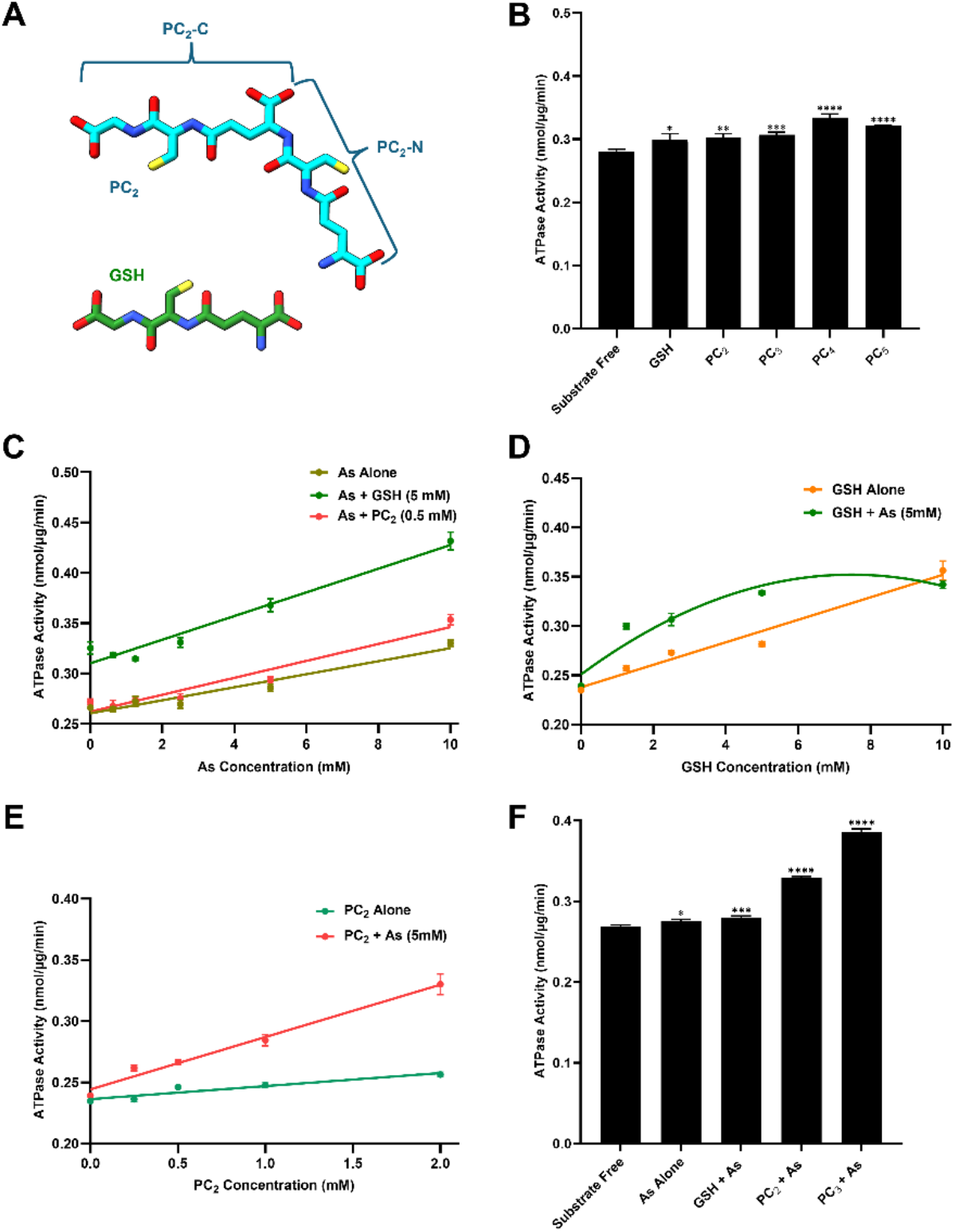
Biochemical characterization of ZmABCC1. (**A**) Chemical structures of GSH and PC_2_ showing the N-terminal (PC_2_-N) and C-terminal (PC_2_-C) glutathione moieties. (**B**) ATPase activity of ZmABCC1 in the presence of GSH and PCs of varying chain lengths at 0.4 mM concentration in the absence of heavy metal. (**C**) ATPase activity of ZmABCC1 as a function of As(III) concentration alone or in the presence of 5 mM GSH or 0.5 mM PC_2_. (**D**) GSH saturation curve of ZmABCC1 ATPase activity in the absence or presence of 5 mM As(III). (**E**) PC_2_ saturation curve of ZmABCC1 ATPase activity in the absence or presence of 5 mM As(III). (**F**) Comparison of ZmABCC1 ATPase activity in the presence of 5 mM As(III) combined with 2 mM GSH, PC_2_, or PC_3_. All data is presented as mean ± standard deviation from triplicates (n=3). Statistical analyses of the data shown in panels B and F were performed by one-way ANOVA followed by Dunnett’s multiple comparison test (ns, P ≥ 0.05 (not statistically significant); *, P < 0.05; **, P < 0.01; ***, P < 0.001; ****, P < 0.0001). Abbreviations: GSH, Glutathione; As, As(III); PC, Phytochelatin.

Here, we characterized the maize ABCC1 homolog, ZmABCC1, and determined that its substrate-stimulated ATPase activity is enhanced by GSH and PCs in complex with As(III). We determined the cryo-EM structures of ZmABCC1 in both its apo state and bound to an As(III)-PC_2_ complex. The substrate-bound structure reveals the binding pose and coordination geometry of the heavy metal complex within substrate binding pocket, identifying key residues responsible for coordinating PC_2_ and the central As(III) atom.

## Methods

### Cloning, expression and purification of maize ABCC1

The maize ABCC1 (ZmABCC1) gene was cloned into pFastBac1 with C-terminal maltose binding protein (MBP) tag and the plasmid was transformed into DH10Bac cells to generate recombinant bacmid plasmid. Baculovirus generation was conducted by introducing the purified plasmid into Sf9 insect cells through transient transfection. Sf9 insect cells at ∼2 x 10^6^ cells/mL cell density in ESF 921 (Expression Systems) insect cell culture media with 2% (v/v) fetal bovine serum (FBS) were infected with P3 baculovirus (2% (v/v)) and incubated at 27 °C for protein expression. After 72 hours, cells were harvested, flash frozen, and stored at -80 °C until use.

ZmABCC1 purification was performed by resuspending the cell pellet in solubilization buffer that included 50 mM Tris-HCl (pH 7.4), 150 mM NaCl, 1% DDM, 0.2% CHS, and protease inhibitors (2 µg/ mL Aprotinin, 10 µM Leupeptin, and 1 µM Pepstatin) and incubated for 1 hour at 4 °C. This was then centrifuged for 1 hour at 17,000 x g at 4 °C and the supernatant was incubated with amylose affinity resin for 1 hour at 4 °C. After the removal of supernatant by gravity flow, resin was washed with a buffer containing 50 mM Tris-HCl (pH 7.4), 150 mM NaCl, 0.05% DDM, and 0.01% CHS and the bound protein was eluted with the same buffer supplemented with 10 mM maltose monohydrate. To remove MBP tag, purified ZmABCC1 protein was incubated with MBP tagged TEV protease (TEV-MBP) in a ratio of 1:5 (TEV protease: ZmABCC1-MBP) overnight at 4 °C. The next day, six subsequent concentration-dilution steps were performed to reduce the maltose concentration in the sample using a 100 kDa concentrator. The sample was then incubated with amylose affinity column for 1 hour to facilitate the binding of the cleaved MBP tag and TEV-MBP protease to the resin. After the incubation, ABCC1 protein was collected in the flow through. Finally, the sample was further purified through size exclusion chromatography (SEC) using Superose 6 gel filtration column with SEC buffer (50 mM Tris-HCl (pH 7.4), 150 mM NaCl, 0.05% DDM, and 0.01% CHS). The peak fractions containing purified ABCC1 were pooled and used for ATPase activity assays. For the cryo-EM sample preparation, the uncleaved ZmABCC1-MBP protein was used, and the final SEC buffer contained 50 mM Tris-HCl (pH 7.4), 150 mM NaCl, 0.06% Digitonin. Then, the peak fractions were collected and concentrated to prepare cryo-EM grids.

### ATPase activity assays

ATPase activity of purified ZmABCC1 was measured by detecting the release of inorganic phosphate through colorimetric method. The hydrolyzed phosphate content was determined by comparing to a standard curve of potassium phosphate (KH_2_PO_4_). The assays were conducted in 96-well plates with a total reaction volume of 50 µL. Each reaction mixture contained 2 µg of purified ABCC1 protein, 5 mM ATP, 6 mM Mg^2+^, 50 mM Tris-HCl (pH 7.4), 150 mM NaCl, 0.05% DDM, and 0.01% CHS. Assays were initiated by adding protein and conducted at 37 °C for 1 hour. After the incubation, reactions were quenched by adding 50 µL of 12% (w/v) sodium dodecyl sulfate (SDS), followed by the addition of 100 µL of a 1: 1 mixture of 12% (w/v) ascorbic acid and 2% (w/v) ammonium molybdate to develop the color. Finally, the absorbance was measured at 850 nm wavelength using a microplate reader.

Each assay was performed in triplicate. Microsoft Excel and GraphPad Prism software were used to analyze data, and presented as the average values ± standard deviation.

### Cryo-EM sample preparation and data collection

Freshly purified ZmABCC1 protein samples were concentrated to 8.3 mg/ mL and 4.5 mg/ mL to prepare grids for apo form and PC_2_-As(III) bound form, respectively. To prepare PC_2_-As(III) bound form grids, 2 mM PC_2_ and 5 mM As(III) were incubated with the protein sample for ∼1 hour on ice. For the grid preparation with both samples, 3 µL of the sample was applied to freshly glow discharged Quantifoil R 1.2/1.3 300 mesh, copper holey carbon grid. Then, the grid was blotted for 3 seconds with 95% humidity at 4 °C using Vitrobot Mark IV (Thermo Fisher) and plunge frozen in liquid ethane.

Data collection for ZmABCC1-apo form grids was done using Glacios electron microscope (Thermo Fisher Scientific), operating at 200kV, equipped with K3 direct electron detector. For PC_2_-As(III) bound ZmABCC1 structure, Titan Krios electron microscope was used for data collection, equipped with a Selectris X energy filter and Falcon 4i camera.

### Cryo-EM data processing and model building

All cryo-EM data processing was performed using CryoSPARC v5. Raw movies were motion-corrected using Patch Motion Correction, and contrast transfer function parameters were estimated using Patch CTF. Movies were manually curated to remove those unsuitable for further processing. Initial templates were generated using Blob Picker followed by several rounds of 2D classification. High-quality 2D class averages were selected as templates for particle picking using Template Picker. Additional rounds of 2D classification were performed, followed by ab initio reconstruction and non-uniform refinement. The resulting 3D volume was used to generate 2D projections, which were subsequently used as templates for a final round of particle picking with Template Picker. Particles were extracted with a box size of 384 pixels and Fourier-cropped to 96 pixels, corresponding to 4-fold binning. All extracted particles were subjected to Ab-initio reconstruction with three classes. The same particle set was then subjected to heterogeneous refinement using the three Ab-initio volumes as initial reference models. Several additional rounds of heterogeneous refinement were performed, after which particles were re-extracted with 1.5-fold binning or without binning. The resulting particles were subjected to non-uniform refinement followed by local refinement. Particles were subsequently subjected to 3D classification without alignment, and particles from the highest-quality class were used for a final round of non-uniform refinement followed by local refinement. Atomic model building and refinement were performed using Coot and Phenix. An AlphaFold model of ZmABCC1 was used as the initial model, and individual domains were manually fit into the cryo-EM density using UCSF ChimeraX. The resulting model was manually adjusted in Coot, followed by real-space refinement in Phenix.

## Results

### Investigation of ZmABCC1 activity towards various chelators and metals

Purification was performed as described previously (Wathukoralage and Culbertson, 2027). Briefly, ZmABCC1 with a C-terminal maltose binding protein (MBP) tag was expressed in Sf9 cells using a baculovirus expression system and purified using amylose resin. Cleavage of the MBP tag was performed with TEV protease, and the cleaved ZmABCC1 was further purified by size-exclusion chromatography (SEC). To investigate the activity of ZmABCC1 toward various thiol chelators such as GSH and PCs, substrate-stimulated ATP hydrolysis (ATPase) assays were performed to determine if the apo-PCs stimulate ZmABCC1 activity. PCs typically occur with varying degrees of oligomerization, most commonly between PC_2_ and PC_5_ (Grill et al., 1987). We measured substrate-stimulated ATPase activity in the presence of 0.4 mM GSH or PCs (PC_2_ through PC_5_), which produced only modest or negligible stimulation of ATPase activity (Figure 1B), indicating that these compounds do not strongly stimulate ZmABCC1 activity in isolation. We next evaluated three proposed metal substrates of AtABCC1, As, Hg, and Cd, to determine whether the metal complex with GSH or PCs influences ZmABCC1 activity. In the case of arsenic (As(III)), addition of arsenite resulted in an apparent increase in phosphate release even in the absence of GSH or PC_2_ (Figure 1C), which was also observed with increasing arsenite in the absence of protein and ATP/Mg^2+^ (Supplemental Figure 1A). Because uncomplexed As(III) is unlikely to be directly transported by ZmABCC1 and a linear increase was observed when protein and ATP were excluded (Supplemental Figure 1A), this increase most likely reflects assay interference rather than true ATPase stimulation. Possible explanations include phosphate contamination in the arsenite stock solution or chemical interference with the molybdate-based color development step of the assay. Despite this background signal, inclusion of 0.5 mM PC_2_ with increasing arsenite produced only modest additional stimulation, primarily at higher arsenite concentrations (i.e., 10 mM). In contrast, when 5 mM GSH was included, the As(III)-dependent increase in ATPase activity displayed a similar or slightly steeper concentration dependence relative to arsenite alone (Figure 1C), suggesting that GSH-As(III) complexes modestly stimulate ZmABCC1 activity. In contrast to arsenite, both Cd(II) and Hg(II) caused a pronounced decrease in ATPase activity at concentrations above ∼1 mM (Supplemental Figure 1B-C). This inhibition is consistent with known biochemical effects of heavy metals. For Cd(II), inhibition may arise from competition with Mg²⁺ at the nucleotide-binding site, whereas Hg(II) likely impairs activity through covalent modification of cysteine residues, resulting in loss of protein function. Notably, inclusion of 5 mM GSH partially restored ATPase activity in the presence of both Cd(II) and Hg(II), and PC_2_ produced a modest protective effect against Cd(II) (Supplemental Figure 1B-C). These observations are consistent with the chelation of heavy metals by GSH or PC_2_, thereby likely reducing the inhibitory impact of these metals on ZmABCC1 by chelating the metals, rather than directly stimulating transport.

We next performed saturation analyses of GSH and PC_2_ in the presence or absence of As(III) (Figure 1D-E). In the absence of As(III), increasing GSH concentration produced a linear increase in ATPase activity. In contrast, inclusion of As(III) resulted in enhanced stimulation at lower GSH concentrations and a saturable response at higher concentrations, indicating that formation of a GSH-As(III) complex increases the apparent affinity of ZmABCC1 relative to GSH alone (Figure 1D). Similarly, PC_2_ titration in the absence of As(III) yielded only a modest, linear increase in ATPase activity. When As(III) was present, PC_2_ produced a pronounced stimulation of ATPase activity across the concentration range tested that increased with rising PC_2_ concentration, supporting the interpretation that the PC_2_-As(III) complex is recognized by ZmABCC1 (Figure 1E). Although a clear saturation plateau was not observed within the tested PC_2_ range (up to 2 mM), this concentration window is narrower than that examined for GSH and does not exclude saturation at higher concentrations. Collectively, these findings support preferential recognition of thiol-coordinated As(III) complexes by ZmABCC1 over the free ligands alone.

Based on the high substrate-stimulated ATPase activity of ZmABCC1 observed in the presence of 2 mM PC_2_ and 5 mM As(III), we next compared GSH and PC_2_ through PC_5_ in the presence of 5 mM As(III). Inclusion of 2 mM GSH increased ATPase only moderately, while adding 2 mM PC_2_ or PC_3_ resulted in progressively larger increase in ATPase, with PC_3_ being higher compared to PC_2_ (Figure 1F). In contrast, assays containing PC_4_ and PC_5_ developed high turbidity upon addition of the assay color development reagents (ascorbic acid and molybdate solutions) at these concentrations, indicating that these complexes precipitate at high concentrations in the presence of As(III). Nonetheless, these data indicate that ZmABCC1 utilizes GSH, PC_2_, and PC_3_ to stimulate transport-associated ATPase activity in complex with As(III), with PC_2_- and PC_3_-As(III) complexes resulting in the highest activity.

### Cryo-EM structure of ZmABCC1

For structure determination, purified ZmABCC1-MBP was prepared for cryo-EM experiments in a buffer containing 0.06% digitonin. The uncleaved MBP-tagged protein was utilized to preserve yield and enable concentration to levels sufficient for generating cryo-EM grids with high particle density. Previous work demonstrated that the MBP tag does not affect ATP hydrolysis kinetic parameters or substrate-stimulated ATPase activity of ZmABCC1 (Wathukoralage and Culbertson, 2027). The final cryo-EM map of ZmABCC1 in the apo form refined to 3.7 Å (Supplemental Figure 2-3), displaying domain organization characteristic of C-family transporters (Figure 2A).

**Figure 2.**
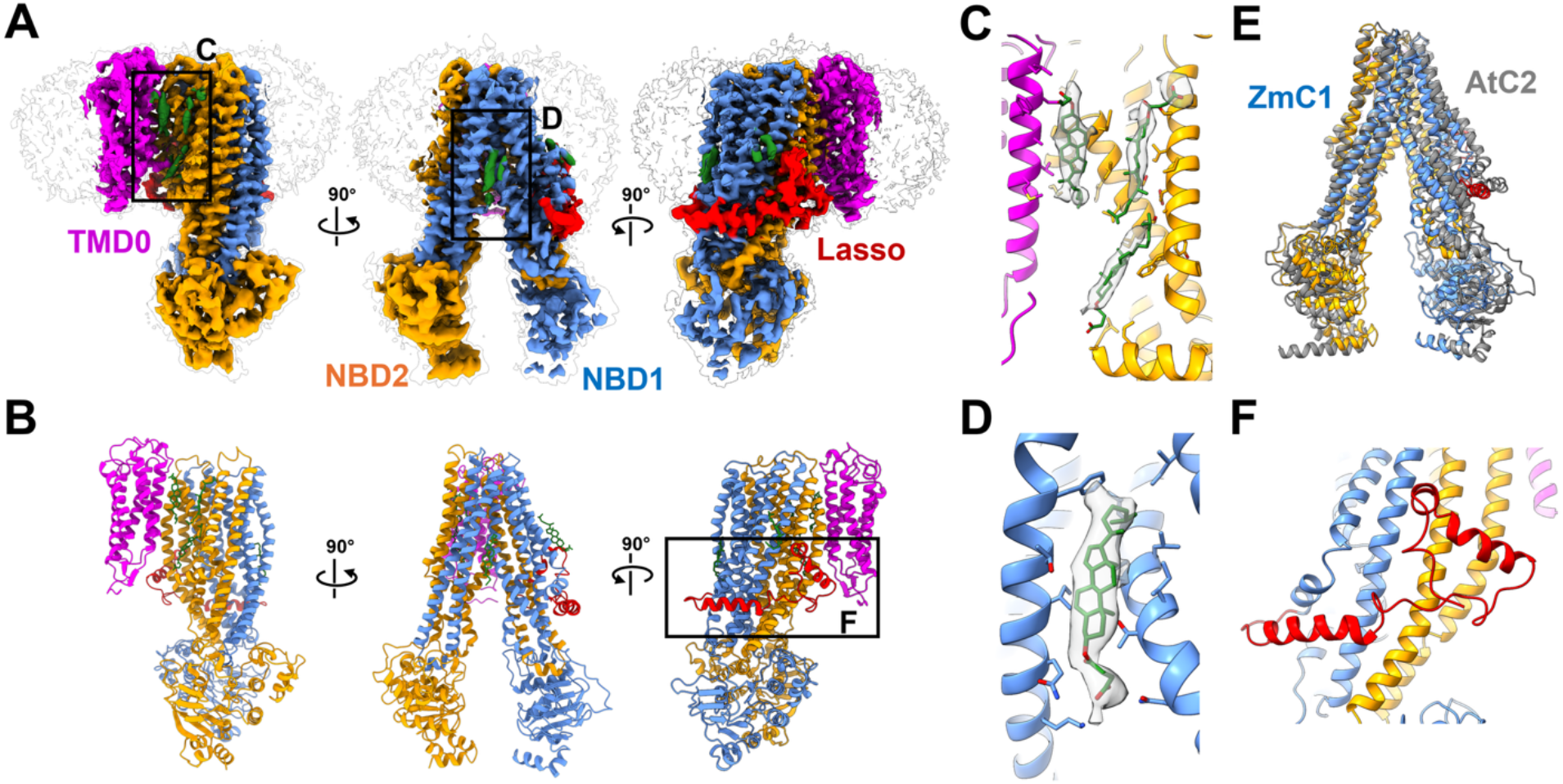
Cryo-EM structure of apo ZmABCC1. (**A**) Orthogonal views of the ZmABCC1 cryo-EM density map. (**B**) Ribbon model of ZmABCC1 shown in orientations identical to panel A. (**C**) Zoomed-in view showing cryo-EM density and fitted models of lipid molecules bridging TMD0 and TMD2. (**D**) Cryo-EM density and modeled cholesteryl hemisuccinate molecule adjacent to TM9 and TM11. (**E**) Structural superposition of apo ZmABCC1 with apo AtABCC2 (grey). (**F**) Zoomed-in view of the lasso domain at the membrane interface. Structural domains of ZmABCC1 are colored as TMD0 in magenta, TMD1-NBD1 in blue, TMD2-NBD2 in orange, and lasso domain in red.

C-family transporters possess three transmembrane domains (TMD0-2), where TMD0 has proposed roles in protein localization or dimerization (Johnson and Chen, 2017; Shinde et al., 2025). In the structure of ZmABCC1, TMD0 contacts TMD2 exclusively (Figure 2A-B), with several interactions mediated by lipid molecules (Figure 2C). The relative positioning of TMD0 in plants differs from that observed in mammals or yeast (Supplemental Figure 4). In plant ABCC1 transporters, including ZmABCC1, TMD0 contacts with TMD2, while in mammalian and yeast homologs, TMD0 contacts with TMD1 (Supplemental Figure 4). Beyond the lipids bridging TMD0 and TMD2, well-defined non-proteinaceous density was observed adjacent to TM9 and TM11 (Figure 2D). This density corresponds in size and shape to a sterol molecule and was modeled as cholesteryl hemisuccinate, which was present in the solubilization and purification buffers (Figure 2D). Comparison of the ZmABCC1 and AtABCC2 apo states reveals closer NBDs in ZmABCC1, with a distance of 25.7 Å between serine residues in the Walker A and Signature motifs in ZmABCC1, compared to 32.3 Å between equivalent residues in AtABCC2 (Figure 2E) (Qiu et al., 2025). The lasso domain, also designated as the L_0_ linker, is an structural motif conserved across C-family ABC transporters (Johnson and Chen, 2017; Khandelwal et al., 2022). The lasso domain is required for proper protein folding, membrane trafficking, and functional activity in C-family transporters (Bakos et al., 1998; Westlake et al., 2003). ZmABCC1 also contains a lasso domain (Figure 2F) and structural overlays demonstrate that the overall conformation of the lasso domain is conserved, showing near identity across mammalian ABCC1, yeast Ycf1, and plant AtABCC2 (Supplemental Figure 5).

### Structure determination of ZmABCC1 bound to PC_2_-As(III) complex

To investigate the binding mode of the PC_2_-As(III) complex to ZmABCC1, we prepared cryo-EM grids of ZmABCC1 in the presence of 2 mM PC_2_ and 5 mM sodium arsenite. Although these concentrations were non-saturating (Figure 1E), preparing higher working concentrations was restricted by challenges in consistently solubilizing PCs at concentrations above 10 mM. Additionally, increasing the volume of substrate stock added to the sample was avoided, as further diluting the protein led to reduced particle density in cryo-EM micrographs. Standard cryo-EM data processing yielded a 2.7 Å map with minimal density for PC_2_-As(III) within the binding cavity (Supplemental Figure 6). Further processing using a mask around the transporter core (TMD1-NBD1-TMD2-NBD2, excluding TMD0) followed by 3D classification without alignment generated a cryo-EM map at 2.9 Å resolution with enhanced density in the substrate-binding site (Figure 3A-C and Supplemental Figure 6-7). This 2.9 Å map with enhanced ligand density exhibited a shorter NBD distance relative to the initial ligand-free 2.7 Å map, consistent with closer NBD distances in substrate-bound conformations reported for other ABC transporters (Johnson and Chen, 2017; Culbertson and Liao, 2025). This alternative processing strategy enhanced the density within the substrate-binding site sufficiently to enable confident model building of the PC_2_-As(III) complex in the ZmABCC1 binding pocket (Figure 3B-C).

**Figure 3.**
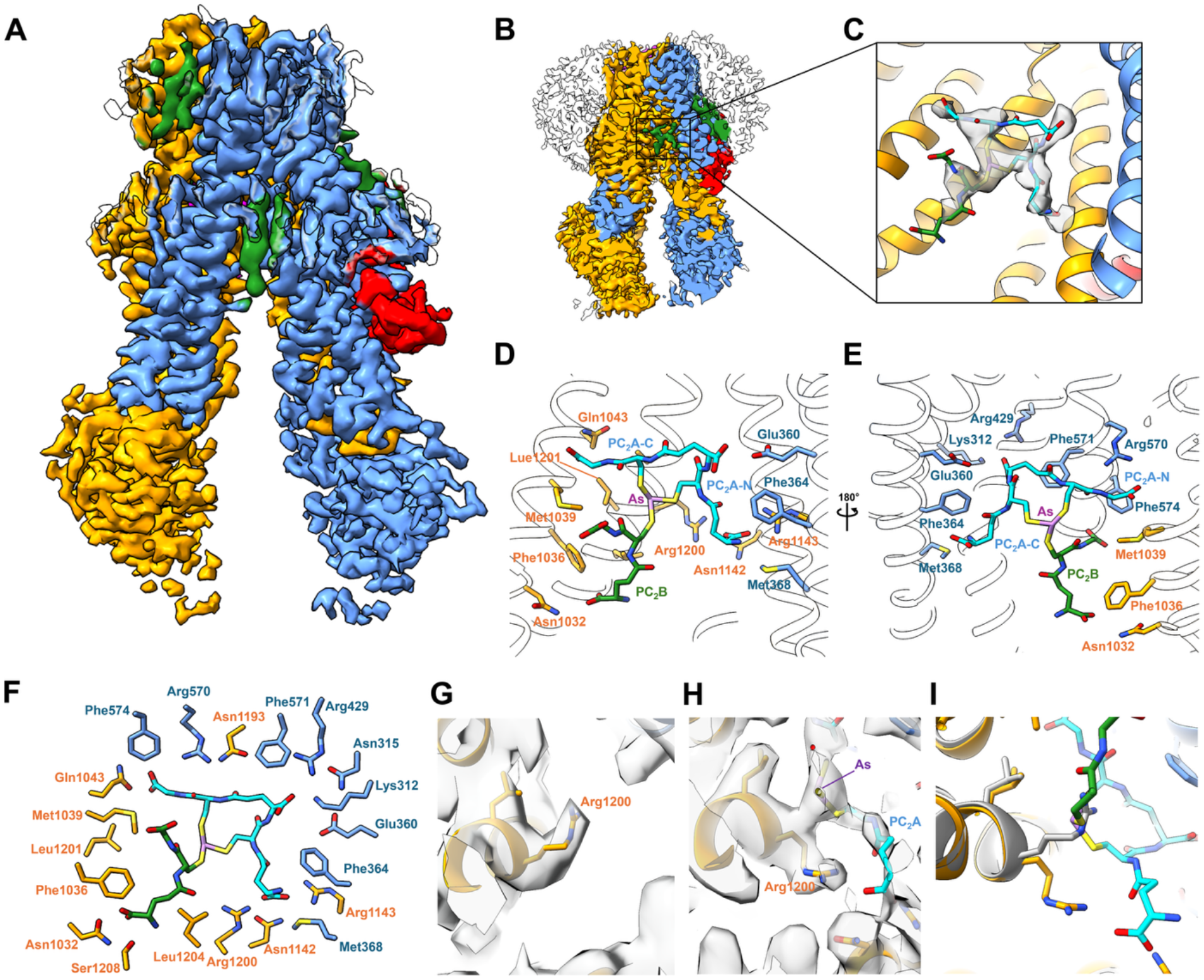
Structure of ZmABCC1 bound to PC_2_-As(III). (**A**) Cryo-EM map of ZmABCC1 bound to PC_2_-As(III). (**B**) Cross-sectional view showing substrate density in green within the central binding cavity. (**C**) Zoomed-in view of the substrate binding cavity with fitted density for the PC_2_-As(III) complex. (**D-E**) Detailed views of the PC_2_-As(III) binding site with interacting amino acid side chains shown as sticks. (**F**) Diagram of residues surrounding the PC_2_-As(III) complex in the binding pocket. (**G-H**) Cryo-EM density and model for Arg1200 in the apo state (**G**) and PC_2_-As(III)-bound state (**H**). (**I**) Overlay of Arg1200 side chain positions in the apo state (grey) and PC_2_-As(III)-bound state (yellow). Abbreviations: As, As(III); PC_2_, Phytochelatin 2.

The PC_2_-As(III) cryo-EM map resolved one full PC_2_ molecule (designated PC_2_A), an As(III) atom, and partial density corresponding to one half of a second PC_2_ molecule (designated PC_2_B). The As(III) atom is coordinated by three sulfur atoms from two distinct PC_2_ ligands, PC_2_A and PC_2_B (Figure 3C-E). PC_2_A contributes two coordinating sulfurs, while PC_2_B provides the third coordinating sulfur. Within the fully modeled PC_2_A ligand, the highest map density corresponds to its N-terminal GSH moiety, designated PC_2_A-N, which occupies the canonical GSH-binding pocket (P-pocket) identified in previously determined ABCC1 structures (Johnson and Chen, 2017; Soong et al., 2025; Sun et al., 2025). The C-terminal portion of PC_2_A, designated PC_2_A-C, extends parallel to the membrane plane and has weaker density relative to PC_2_A-N. The second ligand, PC_2_B, exhibits poor density overall, with resolved features localized primarily around its As(III)-coordinating sulfur atom. Among the components of the (PC_2_)_2_-As(III) complex, the density corresponding to the central As(III) atom exhibits the strongest cryo-EM density, evident from its presence even at higher contour level (Supplemental Figure 8A). At lower contour thresholds, this core density expands in a threefold arrangement (Supplemental Figure 8B-C), consistent with the trigonal pyramidal coordination geometry characteristic of As(III) complexes (Supplemental Figure 8D-G).

Mammalian and yeast ABCC1 homologs possess a conserved GSH-binding site, termed the P-pocket, which contains linear GSH density in a highly similar manner across all available structures (Johnson and Chen, 2017; Soong et al., 2025; Sun et al., 2025). In ZmABCC1, this pocket accommodates the PC_2_A-N moiety and features two cationic sites that coordinate carboxylate groups containing basic residues including Arg1200 and Arg1143 in the first site, and Lys312 and Arg429 in the second (Figure 3D-F). In addition to coordinating a carboxylate group of PC_2_, Arg1200 contacts the non-bonding face of the As(III) atom. In the apo state, Arg1200 is oriented upward, whereas in the PC_2_-As(III)-bound state, it shifts to a downward orientation (Figure 3G-I). In its upward conformation in the apo structure (Figure 3G), the side chain of Arg1200 projects directly into the space occupied by As(III) in the substrate-bound state (Figure 3H-I). Consequently, binding of the (PC_2_)_2_-As(III) complex induces a conformational reorientation of Arg1200 toward the downward position.

The C-terminal GSH moiety, PC_2_A-C, extends horizontally within the binding cavity toward Arg570 in the third cationic binding site (Figure 3D-E). The two cysteine thiolates of PC_2_A converge centrally to occupy two coordination sites of the As(III) center. The third coordination site is occupied by a sulfur atom from the second PC_2_ molecule, PC_2_B, which exhibits poorly defined map density (Figure 3D-F and Supplemental Figure 8A-C). The clearest density in PC_2_B corresponds to the atoms in immediate proximity to As(III), indicating that PC_2_B functions primarily to complete metal coordination rather than establishing extensive contacts with the protein. Nevertheless, the presence of unoccupied volume surrounding PC_2_B suggests that a complete PC_2_ molecule is present within the cavity but the non-As(III) coordinating portion of PC_2_B is largely disordered. As(III) is thus coordinated with trigonal pyramidal geometry by three sulfur atoms provided by two PC_2_ molecules. The non-bonding face of the trigonal pyramidal As(III) center, containing its lone pair, projects toward the aliphatic portion of the side chain of Arg1200. A similar hydrophobic environment surrounding the non-bonding face of As(III) occurs in other protein structures where As(III) is coordinated by three cysteine residues, including an As-binding peptide (PDB ID: 2JGO) (Touw et al., 2007), an As(III)-responsive transcription factor (PDB ID: 6J0E) (Prabaharan et al., 2019), and p53 (PDB ID: 7DHZ) (Chen et al., 2021). In all of these As(III) bound structures, including the ZmABCC1-PC_2_-As(III) structure reported here, As(III) is coordinated by three sulfurs in a trigonal pyramidal geometry, with the non-binding lone pair oriented toward a hydrophobic pocket (Supplemental Figure 8D-G).

### Comparison of PC_2_-As(III) with other substrate-bound ABCC1 structures

As described above, Arg1200 conformationally changes from the upward to downward conformation upon PC_2_-As(III) binding (Figure 3G-I). This upward positioning of Arg1200 in the apo state is enabled by the small side chain of Ala1197, located one turn above Arg1200 on the α-helix, which does not sterically block its upward conformation. In contrast, the corresponding position in mammalian and yeast homologs is occupied by a bulky tryptophan residue (Trp1245 in bovine ABCC1 (bABCC1)). A structural overlay of ZmABCC1-PC_2_-As(III) (Figure 4A) with the bABCC1-LTC4 structure (Johnson and Chen, 2017) (Figure 4B) indicates that Trp1245 in bABCC1 would sterically block the As(III) binding site (Figure 4C-D). This structural difference suggests distinct As(III) binding mechanism between ABCC1 orthologs. In addition, in bABCC1, His335 coordinates a carboxylic acid group within the P-pocket (Figure 4E). In ZmABCC1, this position is substituted by Asn315 (Figure 4F). This loss of basic residue is structurally compensated by a neighboring glutamine-to-arginine substitution (Gln450 in bABCC1 to Arg429 in ZmABCC1) (Figure 4E-F).

**Figure 4.**
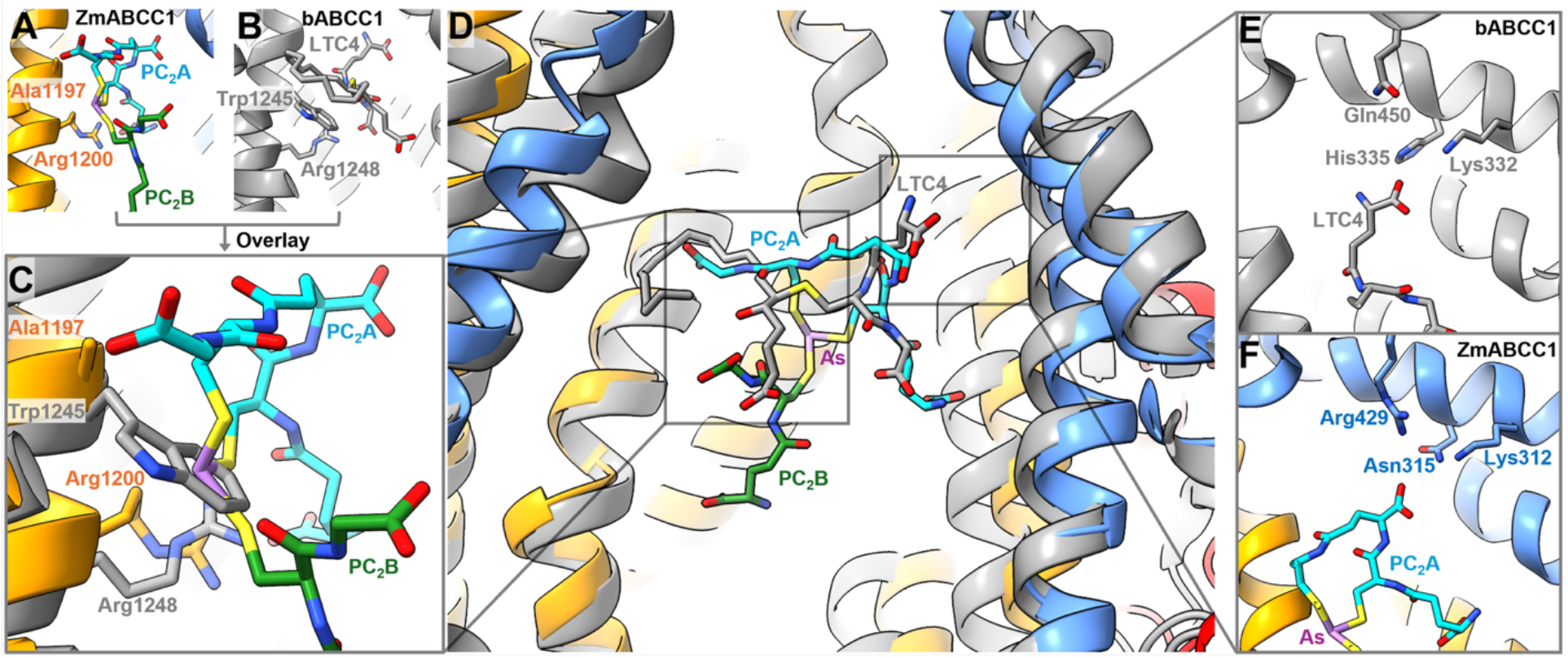
Structural comparison of ZmABCC1 bound to PC_2_-As(III) with bovine ABCC1 (bABCC1) bound to LTC4. (**A**) Substrate binding pocket of ZmABCC1 showing Ala1197 and Arg1200 relative to bound PC2-As(III). (**B**) Substrate binding pocket of bABCC1 bound to LTC4 showing Trp1245 and Arg1248. (**C**) Zoomed-in overlay of panel A and panel B showing the spatial position of Trp1245 in bABCC1 relative to As(III) binding in ZmABCC1. (**D**) Structural superimposition of ZmABCC1 (colored) and bABCC1 (grey) substrate binding cavities. (**E**) Carboxylate coordination pocket of bABCC1 showing side chains His335, Lys332, and Gln450. (**F**) Corresponding carboxylate coordination pocket of ZmABCC1 showing side chains Asn315, Lys312, and Arg429. Abbreviations: As, As(III); PC_2_, Phytochelatin 2; LTC4, Leukotriene C4.

To date, despite strong evidence supporting ABCC1 as heavy metal transporter across plants, mammals, and yeast, no structural information has been reported for these proteins bound to a heavy metal complex. However, cryo-EM structures of Ycf1 and bovine ABCC1 (bABCC1) bound to oxidized glutathione (GSSG) provide structural models for comparing two- or four-coordinate heavy metal complexes, such as cadmium or mercury (Soong et al., 2025; Sun et al., 2025). In all of these structures, substrate recognition relies on a primary anchor point: one glutathione moiety consistently occupies the canonical P-pocket (termed the P1 pocket in Ycf1) in a similar conformation to PC_2_A-N in our PC_2_-As(III) structure (Figure 5) (Johnson and Chen, 2017; Soong et al., 2025; Sun et al., 2025). In contrast, the positioning of the second glutathione moiety in GSSG varies between Ycf1 and bABCC1, and exhibits weaker cryo-EM density in both, suggesting higher structural disorder. GSSG bound to Ycf1 demonstrates that the P-pocket tightly binds the first glutathione to drive selectivity, while the second glutathione moiety occupies a upper pocket designated as P2 (Figure 5B) (Soong et al., 2025). The spatial trajectory of PC_2_A-C in ZmABCC1 occupies a similar region to this upper GSSG moiety in the Ycf1 P2 pocket. Conversely, in the GSSG-bound bABCC1 structure, the second glutathione moiety in GSSG adopts a lower positioning within the plastic binding pocket (Sun et al., 2025). The spatial positioning of the second phytochelatin ligand, PC_2_B, aligns with this lower position observed in bABCC1 (Figure 5C). Thus, while the high-affinity P-pocket anchor is conserved in substrate recognition between orthologs, the secondary ligand moieties, PC_2_A-C and PC_2_B in ZmABCC1, and the second glutathione moiety in GSSG, exhibits structural divergence. Thus, this supports that PC_2_B functions primarily to stabilize and complete the coordination of the central As(III) atom rather than engaging in high-affinity contacts with the transporter substrate-binding pocket.

**Figure 5.**
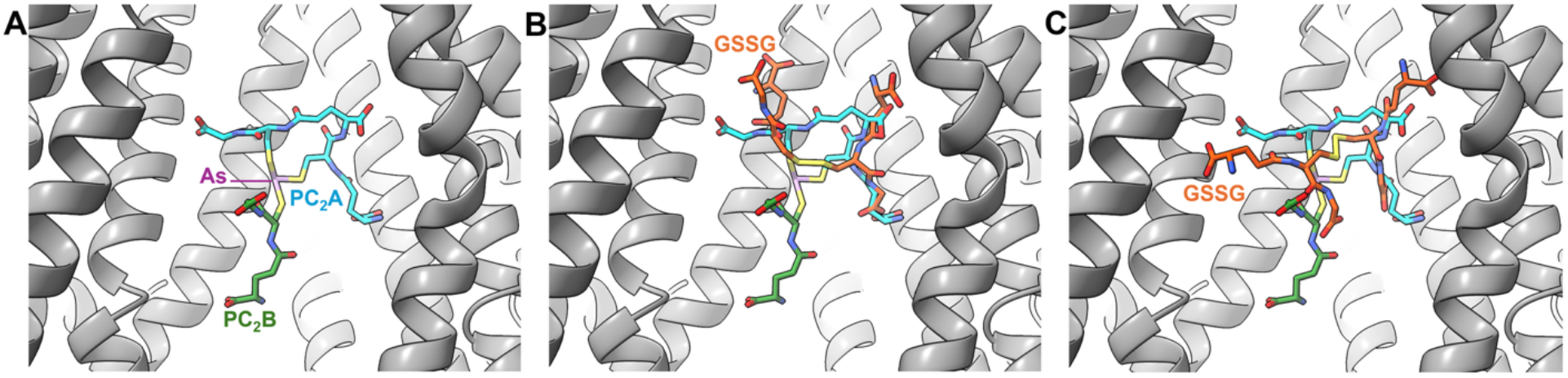
Comparison of ZmABCC1 bound to PC_2_-As(III) with Ycf1 and bABCC1 bound to GSSG. (**A**) Substrate binding cavity of ZmABCC1 bound to PC_2_-As(III) showing PC_2_A (cyan), PC_2_B (green), and As(III) (purple). (**B**) Structural overlay of ZmABCC1 bound to PC_2_-As(III) with Ycf1 bound to GSSG (orange; PDB 9AY0). (**C**) Structural overlay of ZmABCC1 bound to PC_2_-As(III) with bABCC1 bound to GSSG (orange; PDB 9LG9). Abbreviations: As, As(III); PC_2_, Phytochelatin 2; GSSG, Oxidized glutathione.

### Modeling (GSH)_3_-As(III) and PC_3_-As(III)

Based on our in vitro assays, ZmABCC1 utilizes PC_2_ and PC_3_, and GSH to a lesser extend, to stimulate ATPase activity in the presence of As(III) (Figure 1F). This observation is consistent with the threefold coordination requirement of As(III), as each of these thiol chelators can organize to fully satisfy the metal coordination sphere. To evaluate whether the ZmABCC1 substrate-binding pocket can sterically accommodate an (GSH)_3_-As(III) or a PC_3_-As(III) complex, we modeled these potential substrates using our experimentally determined PC_2_-As(III) cryo-EM structure as a guide (Supplemental Figure 9). To construct the (GSH)_3_-As(III) model within the binding pocket (Supplemental Figure 9A), the position of PC_2_B was unchanged because it represents a single GSH moiety. A second GSH molecule was placed to overlay directly with PC_2_A-N within the conserved P-pocket. The third GSH molecule was modeled in a position corresponding to PC_2_A-C, with its glycine residue reoriented upward and toward the adjacent α-helix (Supplemental Figure 9A). Sufficient volume was available in this region to accommodate the reoriented GSH glycine residue without steric clashes, supporting the capacity of ZmABCC1 to accommodate an (GSH)_3_-As(III) complex (Supplemental Figure 9D). To model PC_3_-As(III) (Supplemental Figure 9C), the first two GSH units of the PC_3_ molecule were aligned in identical positions to PC_2_A. The C-terminal segment of PC_3_ was rotated laterally, in which the third cysteine thiolate easily reached an identical coordination position occupied by the PC_2_B sulfur atom (Supplemental Figure 9E). Collectively, this structural modeling indicates that ZmABCC1 can accommodate GSH_3_-As(III), (PC_2_)_2_-As(III), and PC_3_-As(III) complexes without steric hindrance or requiring major conformational changes in the protein backbone.

## Discussion

Thiol-mediated metal detoxification is utilized across all domains of life, including bacteria, plants, animals, and fungi. These systems rely on GSH as the primary thiol source, which, in plants, is synthesized within the plastid and exported to the cytosol (Seregin and Kozhevnikova, 2023). GSH directly chelates several heavy metals and metalloids, including mercury, arsenic, cadmium, and lead. In yeast and mammals, detoxification proceeds via the transport of these GSH-metal complexes into the vacuole or extracellular space (Szczypka et al., 1994; Leslie et al., 2004). In contrast, plants utilize PCs, which are oligomers of glutathione synthesized by phytochelatin synthase (Seregin and Kozhevnikova, 2023). Most plants produce PCs with a degree of polymerization between two and four (PC_2_-PC_4_) (Cobbett and Goldsbrough, 2002; Seregin and Kozhevnikova, 2023). Binding affinity studies indicate that the stability of cadmium-PC complexes increases linearly with chain length up to PC_4_, whereas further polymerization does not enhance complex stability (Jacquart et al., 2017; Wątły et al., 2021). Thus, the elongation of glutathione into phytochelatins increases metal-binding affinity, likely by increasing the local concentration of coordinating thiol groups. While PCs demonstrate higher binding affinity for heavy metals relative to GSH, the evolutionary driver behind PC distribution is complex, as high exposure to toxic metals such as cadmium or arsenic was likely rare in most natural ecosystems (Clemens, 2006). Organisms lacking phytochelatin synthase frequently utilize metallothioneins, which are genetically encoded proteins capable of evolving metal-specific isoforms (Bundy and Kille, 2014). In contrast, non-ribosomally synthesized PCs exhibit identical metal-binding affinities across species because they are identical chemical structures. Because PCs are induced by physiological concentrations of essential micronutrients like zinc and copper, phytochelatin synthase may have originally evolved for basal nutrient homeostasis rather than heavy metal detoxification (Grill et al., 1987; Clemens, 2006). Regardless of whether an organism utilizes GSH or PCs, all rely on an ABCC1 ortholog for vacuolar sequestration or cellular export, though these orthologs exhibit distinct substrate specificities.

The yeast transporter Ycf1 and mammalian ABCC1 utilize GSH to coordinate and export heavy metals including mercury, arsenic, lead, and cadmium (Szczypka et al., 1994; Leslie et al., 2004). In contrast, plant ABCC1 primarily mediates the vacuolar sequestration of heavy metals complexed with PCs (Song et al., 2010; Park et al., 2012). Heterologous expression of plant ABCC1 fails to complement a Ycf1-knockout phenotype in *Saccharomyces cerevisiae* suggests plant ABCC1 does not utilize GSH in similar manner to Ycf1 (Song et al., 2010). However, co-expression of plant ABCC1 with phytochelatin synthase does restore heavy metal resistance in yeast, suggesting plant ABCC1 selectively transports PC-metal complexes over GSH-metal complexes (Song et al., 2010). However, in vitro biochemical characterization of ZmABCC1 described here demonstrates that arsenite stimulated ATPase activity of both GSH and PCs, albeit PC showed higher ATPase compared to GSH (Figure 1). This incremental enhancement indicates that longer PC chain lengths are more efficiently utilized by ZmABCC1, which may be required to reach the threshold of detoxification needed to limit heavy metal toxicity. Beyond metal-coordination preferences (GSH vs PCs), plant ABCC1 orthologs and Ycf1 exhibit distinct heavy metal substrate specificities. Overexpression of Ycf1 in Arabidopsis confers resistance to Pb(II) (Song et al., 2003) whereas knock-out of AtABCC1 and AtABCC2 in Arabidopsis does not alter Pb(II) resistance (Park et al., 2012) suggesting Ycf1 transports Pb(II) while plant ABCC1 do not. Furthermore, Ycf1 displays high transport capacity for GSH-Cd(II) complexes (Soong et al., 2025), whereas plant ABCC1 orthologs transport Cd(II), but prefer As(III) complexes (Song et al., 2014). Thus, although these transporters are broadly classified as thiol-metal efflux pumps, they possess distinct substrate preferences. However, a rigorous evaluation of these functional differences requires direct parallel characterization under identical experimental conditions.

The structural basis for these observed differences in substrate specificity is clarified by the cryo-EM structure of ZmABCC1. A key difference within the substrate-binding pocket occurs at position 1197, which is an alanine in ZmABCC1 but a tryptophan in Ycf1 and mammalian ABCC1 (Figure 4A-B). In ZmABCC1, As(III) contacts the aliphatic portion of the Arg1200 side chain, which would be sterically blocked by the presence of the bulkier tryptophan at position 1197 in yeast and mammalian homologs (Figure 4C). Consistent with a key functional role of this tryptophan residue, mutation of this tryptophan (Trp1225 in Ycf1) to alanine reduces cadmium resistance in *Saccharomyces cerevisiae* (Soong et al., 2025). However, Cd(II) is likely coordinated in a tetrahedral geometry, which remains fully enclosed by coordinating thiol groups, preventing direct structural contact with the protein as observed for As(III). Consequently, this Arg1200-metal interaction is unlikely to occur for Cd(II) or Hg(II) due to their distinct coordination geometries. The presence of alanine at position 1197 may instead allow access to the aliphatic portion of Arg1200, contributing to the preference of plant ABCC1 for As(III) over other metals. Together, these observations indicate that differences in metal coordination geometry and the composition of the substrate-binding pocket likely produce distinct binding modes for GSH-Cd(II) and PC_2_-As(III), which may account for the different metal preferences of plant ABCC1 and Ycf1. A second distinct feature in the binding pocket of plant ABCC1 involves the coordination of the glutathione carboxylate moiety. In yeast and mammalian homologs, a conserved histidine (His335 in bABCC1) coordinates one of the carboxylate groups in the P-pocket (Figure 4E) (Johnson and Chen, 2017; Sun et al., 2025). In plant ABCC1, this position is substituted by an asparagine (Asn315) (Figure 4F). This loss of electrostatic coordination is structurally compensated by a neighboring glutamine-to-arginine substitution, where the arginine side chain (Arg429) extends across the cavity to coordinate the acidic group of PC_2_ (Figure 4F). This localized structural alteration may subtly change the positioning of glutathione within the P-pocket, potentially contributing to the observed differences in substrate specificity.

In addition to GSH and PC_2_, plants also contain higher molecular weight PCs up to PC_4_ or PC_5_ (Meyer et al., 2011). Substrate-stimulated ATPase assays demonstrate that ZmABCC1 activity increases sequentially upon coordination by GSH, to PC_2_, and to PC_3_ (Figure 1F). Modeling supports that the binding pocket accommodates these elongated chain lengths up to PC_3_ (Supplemental Figure 9). However, binding affinity studies show that cadmium complexation increases with PC chain length up to PC_4_ (Jacquart et al., 2017; Wątły et al., 2021). This may reflect the different coordination geometry of Cd(II), which forms a four-coordinate complex, compared with the three-coordinate As(III) complex. Consistent with this distinction, assays with As(III) and either PC_4_ or PC_5_ resulted in high turbidity, suggesting precipitation. This may result from As(III) coordination by three thiol groups, leaving a fourth (and fifth in the case of PC_5_) thiol available to coordinate a second As(III) and form higher-molecular-weight complexes. Further, PCs exceeding PC_3_ are unlikely to further enhance As(III)-stimulated activity, as a single arsenic atom is fully coordinated by PC_3_ (Supplemental Figure 9), leaving terminal thiol segments in PC_4_ or greater disordered within the cavity. Thus, PC_3_ likely represents the optimal PC chain length for As(III) coordination and transport by ZmABCC1, as longer PCs may form higher molecular weight complexes and leaving terminal thiol segments disordered in the binding pocket.

Cryo-EM structures of yeast and bovine ABCC1 bound to GSSG, along with human ABCB6 bound to GSH and PC_2_ coordinated Cd(II), provide insights into substrate binding geometry. Across these GSSG-bound structures and the ZmABCC1-PC_2_-As(III) complex, the glutathione moiety bound in the P-pocket exhibits the strongest density, whereas the remainder of the ligand is less defined (Soong et al., 2025; Sun et al., 2025). Extending out from the P-pocket, the density is weaker in the GSSG structures and there is no consistent organization. Both the GSSG-bound structures show no high-affinity contact with the second GS not bound to P-pocket and neither interact with the aliphatic backbone like the non-polar lone pair of As(III) does in the ZmABCC1-PC_2_-As(III) structure (Figure 3H). In addition, the structure of human ABCB6 was also resolved bound to (GSH)_2_-Cd(II) and (PC_2_)_2_-Cd(II). However, in contrast to ABCC1, ABCB6 is a homodimer with two-fold symmetry and coordinates GSH or PC_2_ molecules from opposing subunits, resulting in a stable substrate-bound state with strong density for the Cd(II) atom (Choi et al., 2024). Thus, ABCC1 and ABCB6 employ distinct binding mechanisms. ABCC1 stably binds a single GSH moiety while utilizing a flexible, non-specific hydrophobic pocket for the remainder of the molecule, whereas ABCB6 forms direct electrostatic interactions from both ends of the ligand to stabilize the complex through conserved residues.

## Conclusion

In summary, this work provides the structural basis for PC-mediated heavy metal sequestration by a plant ABCC1 transporter. Structural determination of ZmABCC1 in complex with an PC_2_-As(III) complex reveals how two distinct PC ligands converge to satisfy the trigonal pyramidal coordination geometry of As(III), positioning the N-terminal glutathione moiety of a PC molecule within a conserved P-pocket. Comparing ABCC1 with different orthologs reveals key structural differences, including an Ala-to-Trp substitution at position 1197 and differences in cation binding site, that distinguish plant ABCC1 from yeast and mammalian homologs. Biochemical assays confirm that while ZmABCC1 accommodates multiple thiol chelators, increasing PC chain length increases ATPase activity. These insights define the core principles for heavy metal recognition in plant vacuolar transporters, and may aid in engineering crop varieties with enhanced metal tolerance and reduced toxic accumulation in edible tissues.

## Data availability

The cryo-EM density maps generated in this study have been deposited to EMDataBank under accession codes 79022 (apo) and 79021 (PC_2_-As(III) bound). The atomic coordinates have been deposited in the Protein Data Bank under accession codes 380Y (apo) and 380X (PC_2_-As(III) bound). Accession codes for both cryo-EM maps and atomic coordinates are summarized in Supplementary Table 1.

## Acknowledgements

Research reported in this publication was supported by the National Institute of General Medical Sciences of the National Institutes of Health under Award Number R35GM166048. The content is solely the responsibility of the authors and does not necessarily represent the official views of the National Institutes of Health. Cryo-EM datasets were collected at Iowa State University Cryo-EM facility with assistance from Jyoti Gupta and at the Michigan State University Cryo-EM facility by Sundharraman Subramanian.

